# The Contributions of Eye Gaze Fixations and Target-Lure Similarity to Behavioral and fMRI Indices of Pattern Separation and Pattern Completion

**DOI:** 10.1101/2021.11.20.469401

**Authors:** Daniel K. Bjornn, Julie Van, C. Brock Kirwan

**Affiliations:** Department of Psychology, Brigham Young University; Neuroscience Center, Brigham Young University

**Author notes:** Corresponding Author: Brock Kirwan, Ph.D.

**Keywords:** fMRI, Memory, Eye Movements, Hippocampus, Mnemonic Discrimination, Pattern Completion

## Abstract

Pattern separation and pattern completion are generally studied in humans using mnemonic discrimination tasks such as the Mnemonic Similarity Task (MST) where participants identify similar lures and repeated items from a series of images. Failures to correctly discriminate lures are thought to reflect a failure of pattern separation and a propensity toward pattern completion. Recent research has challenged this perspective, suggesting that poor encoding rather than pattern completion accounts for the occurrence of false alarm responses to similar lures. In two experiments, participants completed a continuous recognition task version of the MST while eye movement (Experiments 1 and 2) and fMRI data (Experiment 2) were collected. In Experiment 1, we replicated the result that fixation counts at study predicted accuracy on lure trials (consistent with poor encoding predicting mnemonic discrimination performance), but this effect was not observed in our fMRI task. In both experiments we found that target-lure similarity was a strong predictor of accuracy on lure trials. Further, we found that fMRI activation changes in the hippocampus were significantly correlated with the number of fixations at study for correct but not incorrect mnemonic discrimination judgments when controlling for target-lure similarity. Our findings indicate that while eye movements during encoding predict subsequent hippocampal activation changes for correct mnemonic discriminations, the predictive power of eye movements for activation changes for incorrect mnemonic discrimination trials was modest at best.

---

Pattern separation and pattern completion are computational processes involved in mnemonic discrimination, or the discrimination of very similar memories. Pattern separation allows for similar memory traces to be encoded as dissimilar as possible, ensuring that distinct memories remain distinct (Yassa and Stark, 2011; Hunsaker and Kesner, 2013). In contrast, pattern completion is the process whereby a partial or degraded cue leads to the retrieval of a memory trace (Hunsaker and Kesner, 2013). In extreme cases, pattern completion may lead to catastrophic interference, as all similar memories may be retrieved given a single retrieval cue (Norman and O’Reilly, 2003).

The Mnemonic Similarity Task (MST) is a behavioral mnemonic discrimination paradigm that taxes pattern separation processes in human participants. In the MST, participants review images and indicate whether each item has been seen before (Stark et al., 2019). Some images are repeats of items presented before (targets), whereas others are very similar but not the same (lures). Mnemonic discrimination performance in the MST relies on hippocampal-dependent pattern separation processes (Kirwan et al., 2012); however, it is unclear if unsuccessful mnemonic discrimination is due to pattern completion or to another process such as poor encoding. If pattern separation is successful, then a high-fidelity representation of a target stimulus should be strongly encoded, which in turn, will be available to guide a mnemonic discrimination decision when presented with a similar lure. However, as the similarity between the target and lure increases, discrimination performance will be driven more by pattern completion processes (Anderson et al., 2016). According to computational models, this trade-off between pattern separation and pattern completion is driven by attractor dynamics and thus is influenced by the strength or sharpness of the original memory encoding (Knierim and Neunuebel, 2016). Accordingly, mnemonic discrimination is influenced by both the quality of encoding or the strength of the initial memory representation and the degree of similarity between the target and lure stimulus.

In a previous study, eye tracking was used as an implicit measure of memory encoding strength while participants completed the MST (Molitor et al., 2014). Lure items incorrectly identified as “old” (lure false alarms or LFAs) had fewer fixations during the study episode than lure items that were correctly identified as “similar” (lure correct rejections or LCRs), indicating that poor encoding or failure of encoding may account for false alarm responses to lure items. Other recent work monitoring eye movement in partial-cue memory tasks has since challenged this finding (Wynn et al., 2020). Participants who were shown a degraded novel cue tended to focus on regions looked at when viewing similar items. Reinstatement of encoding eye movements was negatively associated with the accuracy of lure responses, indicating that the retrieval of existing representations, or pattern completion, contributed to LFAs (Wynn et al., 2020).

Previous research has demonstrated a relationship between eye movement fixations at encoding and subsequent hippocampal activation such that more fixations at study predict a larger decrease in activation (or repetition suppression) in the hippocampus at retrieval (Liu et al., 2017). While more fixations are associated with stronger repetition suppression, it is unclear if this effect is modulated by target-lure similarity as this factor was not manipulated in the previous study.

Here, we present two experiments replicating the methods of previous studies while accounting for factors that may affect memory performance, including visual complexity of the stimuli and target-lure similarity in order to examine how memory encoding strength (as indexed by eye fixations during study) and target-lure similarity drive behavioral responses and neural activation changes during mnemonic discrimination. We hypothesized that stronger initial encoding as indicated by more eye fixations would predict better subsequent mnemonic discrimination performance. We also hypothesized that target-lure similarity would be a strong predictor of behavioral performance. Finally, consistent with previous fMRI literature, we predicted that more fixations at encoding would be associated with greater repetition suppression in the hippocampus at retrieval.

## Materials and Methods

### Participants

Experiment 1 replicated the behavioral and eye tracking methods of Molitor et al. (2014). Accordingly, we conducted an a priori power analysis using the effect sizes reported in the previous study. Using the lowest reported effect size in the previous paper (η_p_^2^ = 0.17 for the main effect of presentation in a repeated measures ANOVA), 60 participants were required for sufficient statistical power at the 0.80 level. Allowing for attrition, a total of 67 participants (32 male, 35 female) were recruited from the university community for Experiment 1. Data were excluded from four participants in Experiment 1 due to technical issues with the eye tracker (n=3) and uncorrected vision (n=1). After exclusions, a remaining total of 63 participants (29 male, 34 female) were included for data analysis in Experiment 1, mean age 20.0 years (*SD* = 1.91).

Experiment 2 was conducted in the MRI scanner while fMRI and eye tracking data were collected. We recruited 49 right-handed native English-speaking individuals (22 male, 27 female) to participate in Experiment 2. To ensure the safety of those being scanned, we excluded participants with any contraindications for MRI. We excluded data from 14 individuals. Four participants were removed from further analysis given challenges related to the eye tracker calibration. Data from six other participants were not included due to one or more of the following: technical difficulties with the scanner, having previously participated in an experiment using the same stimulus set, safety considerations, not adhering to task instructions, or time constraints preventing the completion of the experiment. Lastly, data from four participants were excluded for excessive motion in the scanner. The limit for excessive motion was set at 1.5 times the interquartile range above the 75^th^ percentile of the number of TRs lost for subjects. After exclusions, we analyzed a remaining total of 35 participants (18 male, 17 female), mean age 22.06 years (*SD* = 2.41).

All participants had normal or corrected-to-normal vision, normal color vision, and reported no history of traumatic brain injury, learning disability, or psychological or neurological diagnosis. We offered course credit to compensate for participation in Experiment 1. Participants in Experiment 2 had the option of monetary compensation or a ¼-scale 3D-printed model of their brain. All protocols were approved by the Brigham Young University Institutional Review Board and all participants provided written, informed consent before participating in the study.

### Stimulus Materials

Stimuli for both experiments were taken from the Mnemonic Similarity Task (Stark et al., 2019) and consisted of color photographs of individual objects on a white background. Stimuli were grouped into five different types, namely subsequent repeat, subsequent lure, repeat, lure, and foil (see Figure 1). Target-lure similarity was normatively rated in a separate behavioral experiment (see Anderson et al., 2016) where an independent sample of participants rated the similarity of target-lure pairs on a 7-point Likert scale. Ratings were subjective and continuous, though selecting the most and least similar pairs resulted in a discontinuity. The current experiment utilized 90 target-lure pairs with the highest and 90 pairs with the lowest similarity ratings from the larger set of 600 target-lure pairs used in our previous publication (Anderson et al., 2016). Each stimulus was assessed for visual complexity by calculating the feature congestion measure of visual complexity for each image (https://dspace.mit.edu/handle/1721.1/37593; Rosenholtz, Li, & Nakano, 2007).

**Figure 1.**
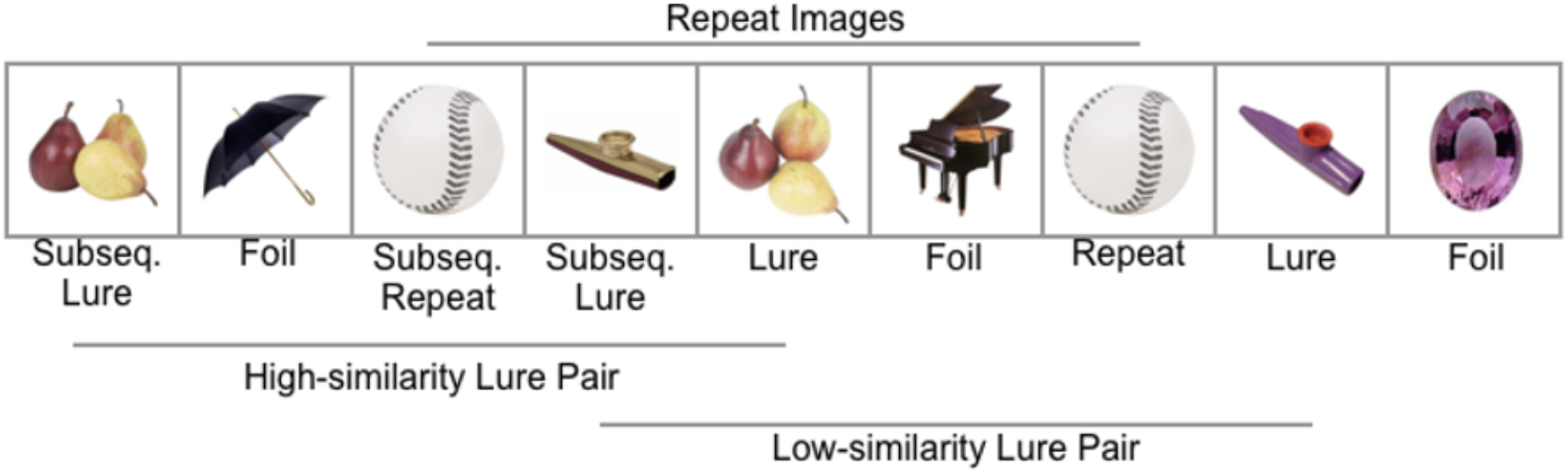
Sample sequence of stimuli. Stimuli were presented one at a time on the center of the screen in a pseudo-random order. Repeat and lure trials were presented with lag times of 4, 12, and 40 trials between first and second presentations. Lag times of 4 trials are presented for simplicity.

### Apparatus

#### Eye Trackers

In Experiment 1, stimuli were presented on a BenQ monitor with a resolution of 1600 × 900 pixels and a refresh rate of 60 Hz (BenQ Corporation, Taipei, Taiwan). Eye movement data were collected using an EyeLink 1000 Plus eye tracker sampled at 1000 Hz from the right eye (SR Research, Ontario, Canada). The eye tracker connected to a host computer running EyeLink 5.0 tracking software. We used Experiment Builder software to program the experiment and collected participant responses using a ResponsePixx Handheld button box (SR Research, Ontario, Canada; VPixx Technologies, Quebec, Canada).

In Experiment 2, stimuli were presented using an MRI compatible BOLDscreen monitor with a resolution of 1920 × 1200 pixels and a refresh rate of 60 Hz (Cambridge Research Systems Ltd., Rochester, United Kingdom). Eye movement data were collected on an MRI compatible EyeLink 1000 Plus long-range eye tracker with a spatial resolution of 0.01° and sampled at 1000 Hz from the right eye (SR Research, Ontario, Canada). We ran EyeLink 5.0 tracking software on a host computer connected to the eye tracker, programmed the experiment with PsychoPy (version 1.85.6; Peirce, 2007, 2008), and integrated the eye tracker using the Eyelink Developers Toolkit (SR Research, Ontario, Canada). Participant responses were collected using a 4-button cylinder MRI compatible button box (Cambridge Research Systems Ltd, Rochester, United Kingdom).

In both experiments, a 9-point eye-tracking calibration was performed before each block for accurate data collection. Successful calibration required an average error of less than 0.5° and a maximum error of less than 1.0°.

#### MRI Scanner and Parameters

MRI scans were conducted on a Siemens 3T Tim Trio scanner at the Brigham Young University MRI Research Facility. We acquired a structural scan from each subject for functional localization using a T1-weighted magnetization-prepared rapid acquisition with gradient echo (MP-RAGE) sequence with the following parameters: TR = 1900 ms; TE = 2.26 ms; number of slices = 176; slice thickness = 1 mm; matrix size = 256 × 215; field of view = 256 × 215 mm; voxel size = 1 mm^3^; flip angle = 9°. A high resolution T2-weighted structural scan centered on the medial temporal lobe and aligned to the long axis of the hippocampus was also obtained using the following parameters: TR = 6260 ms; TE = 64 ms; 19 interleaved slices; slice thickness = 3 mm; matrix size = 512 × 512; field of view = 200 mm^2^; voxel size = 0.39 × 0.39 × 3 mm; flip angle = 178°; averages = 2. Functional scans were collected using a gradient-echo, echo-planar, T2*-weighted pulse sequence with the following parameters: TR = 1900 ms; TE = 28 ms; number of slices = 29 interleaved; slice thickness = 1.8 mm; matrix size = 128 × 128; field of view = 230.4 mm^2^; voxel size = 1.8 mm^3^; flip angle = 90°. Slices were positioned parallel to the long axis of the hippocampus before acquisition.

#### Experiment 1: Eye Tracking Replication Study

Participants performed four blocks of a continuous recognition memory task with images of everyday objects while eye movement data were collected. Images were presented one at a time while participants responded to each trial by indicating whether the presented item was new (i.e., something they had never seen in the experiment), similar (i.e., something like an item they had seen before, but not exactly the same), or old (i.e., an exact repeat of something they had seen before). Items only repeated within a block and no items carried over to another block. Accordingly, stimuli were grouped into five different types: subsequent repeat, subsequent lure, repeat, lure, and foil (see Figure 1). Subsequent repeat and repeat items were identical in each respective pair. Subsequent lure and lure items were similar images of the same type of object, but not exact repeats (e.g., a light brown teddy bear and a dark brown teddy bear). Foil items were images presented only once in the task.

Based on procedures used in the previous study, each trial began with a drift check that also served as the inter-trial interval (Molitor et al., 2014). For the drift check, participants looked at a circle presented in the middle of the screen and pressed a button to ensure that eye position was still correctly calibrated. We performed a calibration before proceeding to the next trial for drift checks that indicated the eye position was beyond acceptable measures. Following the drift check, images were presented for 2.5 seconds, followed by an answer period with response options of “old”, “similar”, and “new” displayed on the screen. Participants were instructed to respond only during the response period, which was terminated with a button press. Images were sized to 17.5° of visual angle on its longest axis and presented at the center of the screen with a gray background. Each task block included 116 trials, 44 of which were foil presentations, 18 subsequent repeat presentations, 18 subsequent lure presentations, 18 repeat presentations, and 18 lure presentations. We grouped lure and repeat presentations into different lag delays of 4, 12, and 40 trials with six trials of each stimulus type falling in each lag delay.

Lure pairs were grouped evenly between high and low-similarity pairs. We integrated these pairs into the run, such that out of the 18 lure pairs presented in each run, nine were high-similarity pairs and nine were low-similarity pairs. Furthermore, half of the trials in each lag were high-similarity and half were low-similarity.

#### Experiment 2: fMRI with Eye Tracking

In Experiment 2, we integrate eye movement data with fMRI to examine activity in the hippocampus as a function of the number of fixations made on an item. This approach enables us to understand how fixation counts affect hippocampal activation and whether this effect changes between LCRs and LFAs. Further, it allows us to observe whether hippocampal activity for repeated images changes as a function of the number of fixations on the first presentation of the stimulus (Liu et al., 2017) in the context of a difficult mnemonic discrimination task.

Stimuli and procedures for Experiment 2 followed those in Experiment 1 with the following exceptions: to ensure an adequate number of trials for fMRI analysis, participants completed five blocks of 116 trials rather than four blocks. Trial timing was also held consistent for fMRI analysis by excluding drift checks that preceded each trial. The answer period was held at 1.5 seconds and not self-paced to maintain trial timing as well.

### Data Processing

#### Behavioral Data

Behavioral data were corrected for response bias prior to analysis. Corrected recognition scores were calculated by subtracting the proportion of “old” responses to foil trials from the proportion of “old” responses to repeat trials. Lure discrimination indices (LDI) were calculated by subtracting the proportion of “similar” responses to foil trials from the proportion of “similar” responses to lure.

#### Eye Movement Data

Raw eye tracking data were processed using Data Viewer software (SR Research, Ontario, Canada). Saccades were defined as movement of at least 0.1° with velocity equal to or greater than 30°/sec and acceleration of 8000°/sec^2^. Fixations were defined as periods of time between saccades that were not part of blinks. Saccades and fixations were removed from analysis if they were outside the maximum stimulus bounds (17.5° × 17.5° visual angle) or immediately preceding or following a blink. Fixations less than 50 ms or greater than 1500 ms were also removed from analysis (Hannula et al., 2010; Carter and Luke, 2020). This resulted in the loss of 3.3% of the data and 0.21% of the data, respectively. Memory-based fixation counts were calculated by subtracting the number of fixations for correct foil trials (i.e., correct rejections) from all other trial types (Molitor et al., 2014). Thus, positive scores denote more fixations than correct foil trials and negative scores denote fewer fixations than correct foil trials. In all analyses, outliers were fenced to 1.5 times the interquartile range above or below the 75^th^ and 25^th^ percentiles, respectively.

#### MRI Data

We analyzed MRI data using dcm2niix (version v1.0.20190902; Li et al., 2016) and Analysis of Functional Neuroimaging (AFNI; version AFNI_18.3.16; Cox, 1996). All MRI data are available at https://openneuro.org/datasets/ds003711/ and analysis scripts are available at https://github.com/Kirwanlab/mst_eyetracking. MRI data were first converted from DICOM to NIfTI format and de-faced to preserve anonymity. Functional scans were not slice-time corrected. Motion correction for each functional run was calculated relative to the acquisition volume (TR) with the smallest number of outlier voxel values. Structural scans were coregistered with this TR as well. Rotated structural scans were skull-stripped and spatially normalized to the MNI template using a nonlinear diffeomorphic transformation. The motioncorrection and MNI-normalization spatial transformations for the functional data were concatenated and applied in a single step to reduce the number of interpolations and thus the blurring of the functional data (Polimeni et al., 2018). We maintained the spatial resolution of the functional scans at 1.8mm^3^ and no additional blurring was done as part of the preprocessing. Functional data were scaled by the mean of the functional run. Volumes with motion events were censored from further analysis if the derivative of the motion parameters had a Euclidean Norm>0.3 (https://afni.nimh.nih.gov/pub/dist/doc/program_help/1d_tool.py.html). Coverage masks were calculated for each participant and each run such that voxels with very low EPI signal were masked and excluded from further analysis.

A single subject regression model was created for each participant that included regressors for motion (6 regressors per scan run) and polynomial regressors accounting for scanner drift. Behavioral regressors in the model coded for subsequent hits, hits, subsequent LCRs with high similarity lures, LCRs with high similarity lures, subsequent LCRs with low similarity lures, LCRs with low similarity lures, subsequent LFAs with high similarity lures, LFAs with high similarity lures, and all other trials. Note that there were too few false alarms to low-similarity lures to reliably model that response type separately. Trial onsets for the conditions of interest were convolved with a square wave of 2.5 s duration and a canonical hemodynamic response function. We set correct foil trials as the model baseline. The resulting β-coefficients were then entered into group-level analyses. A gray matter mask was created based on the MNI template and was combined with the coverage masks from each participant to create a group-level mask. Note that due to coverage constraints, we did not obtain full brain coverage. To identify regions associated with memory performance, we contrasted the subsequent hit and hit conditions using a 2-tailed t-test, which identified brain regions that exhibited repetition suppression and repetition enhancement. Significant clusters of activation were determined using AFNI’s equitable thresholding and clustering (ETAC) function (Cox, 2019), which uses a combination of voxel-wise and spatial extent thresholds to maintain a false-positive rate (FPR) of 5%. Repetition suppression effects were then investigated within these functionally-defined regions of interest (ROIs) by extracting the mean difference in parameter estimates (i.e., betas) between encoding and retrieval for each behavioral outcome (hits, LFAs, LCRs) and then calculating the Pearson correlations between difference scores and mean fixations. This approach allowed us to avoid “double dipping” in our data to avoid spuriously large correlations (Vul et al., 2009).

## Results

### Behavioral Data

Participants performed well across both experiments, correctly identifying 89%-94% of targets and 55%-60% of lures. Proportions of responses can be found in Table 1. We observed significantly better performance for target recognition (corrected recognition scores) than for lure discrimination (LDIs collapsing across lure similarity levels) for Experiment 1, *t*(62) = 24.01, *p* < .001, 95% CI [0.34, 0.40] and Experiment 2, *t*(34) = 14.05, *p* < .001, 95% CI [0.24, 0.32], indicating that participants were better at recognizing repeat items than they were at discriminating similar lures.

**Table 1.** Proportion of Responses by Trial Type and Response. *Note: SD* = standard deviation, First = subsequent repeats and lures, Foil = items only presented once, Lure = similar items, Repeat = items presented twice

| Trial Type | Similarity | Response | Mean (SD) Proportion Responses |  |
| --- | --- | --- | --- | --- |
|  |  |  | Exp. 1 | Exp. 2 |
| Foil | NA | “old” | 0.01 (0.01) | 0.00 (0.01) |
|  |  | “similar” | 0.04 (0.03) | 0.02 (0.03) |
|  |  | “new” | 0.96 (0.03) | 0.97 (0.03) |
| First | NA | “old” | 0.00 (0.01) | 0.00 (0.01) |
|  |  | “similar” | 0.04 (0.03) | 0.02 (0.02) |
|  |  | “new” | 0.96 (0.03) | 0.98 (0.02) |
| Target | NA | “old” | 0.94 (0.05) | 0.89 (0.08) |
|  |  | “similar” | 0.04 (0.03) | 0.08 (0.05) |
|  |  | “new” | 0.02 (0.04) | 0.04 (0.04) |
| Lure | Low | “old” | 0.06 (0.06) | 0.06 (0.05) |
|  |  | “similar” | 0.83 (0.12) | 0.77 (0.17) |
|  |  | “new” | 0.12 (0.09) | 0.17 (0.17) |
| Lure | High | “old” | 0.59 (0.18) | 0.47 (0.16) |
|  |  | “similar” | 0.38 (0.18) | 0.48 (0.16) |
|  |  | “new” | 0.03 (0.04) | 0.05 (0.06) |

To examine the effect of target-lure similarity on lure discrimination, we subjected LDI scores (Table 2) to a repeated-measures ANOVA with similarity (high, low) as a within-subjects factor and experiment (Experiment 1, Experiment 2) as a between-subjects factor. We observed a strong main effect of similarity, *F*(1,96) = 320.29, p<.001, η_p_^2^ = .77, and a similarity × experiment interaction, *F*(1,96) = 14.98, p < .001, η_p_^2^ = .14. The main effect of experiment was not significant, *F*(1,96) = 2.34, p = .13, η_p_^2^ = .02.

**Table 2.**
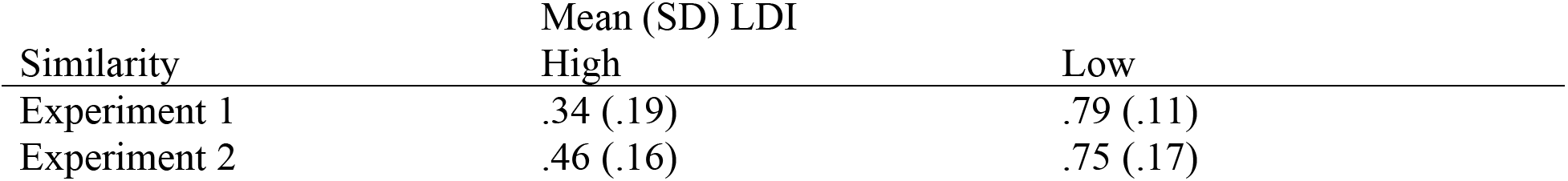
Lure Discrimination Index (SD) for High- and Low-Similarity Lures by Experiment. *Note: SD =* standard deviation.

| Similarity | Mean (SD) LDI |  |
| --- | --- | --- |
|  | High | Low |
| Experiment 1 | .34 (.19) | .79 (.11) |
| Experiment 2 | .46 (.16) | .75 (.17) |

### Experiment 1: Eye Movement Data

To replicate the analysis of Molitor et al. (2014), we entered memory-based fixation counts into a repeated-measures ANOVA with factors coding for presentation (1st, 2nd) and response type (hit, LCR, LFA) collapsing over high- and low-similarity levels. There were significant main effects for presentation, *F*(1, 62) = 9.04, p < .01, η_p_^2^ = .13, and response type, *F*(2, 124) = 32.30, p < .001, η_p_^2^ = .34. The interaction between presentation and response type was also significant, *F*(2, 124) = 11.39, p < .001, η_p_^2^ = .16. There were more fixations on the first presentation of subsequent LCRs than fixations on the first presentation of subsequent LFAs, *t*(62) = 4.37, p < .001, 95% CI [0.15, 0.41], thus replicating findings of Molitor et al. (2014). However, we also observed more fixations on the second presentation of LCRs than fixations on the second presentation of LFAs, *t*(62) = 3.32, p < .01, 95% CI [0.07, 0.30].

We next examined the effects of similarity on memory-based fixations for lures using a repeated measures ANOVA with factors for presentation number (first, second), response type (LCR, LFA), and similarity rating (high, low). We found significant main effects for presentation, *F*(1,46) = 19.14, p < .001, η_p_^2^ = .31, and outcome, *F*(1,46) = 9.36, p = .004, η_p_^2^ = .17, and a presentation × response type × similarity interaction, *F*(1,46) = 4.03, p =.01, η_p_^2^ = .14. Post-hoc comparisons revealed that there were more memory-based fixations during the first presentation for subsequent LCRs than LFAs for both low-similarity lures, *t*(46) = 2.60, p = .013, 95% CI [0.15, 1.21], and high-similarity lures, *t*(62) = 2.86, p = .006, 95% CI [0.08, 0.46]. During the second presentation there were significantly more memory-based fixations for high-similarity LCRs than either high-similarity LFAs, *t*(62) = 3.66, p<.001, 95% CI [0.15, 0.52], or low-similarity LCRs, *t*(62) = 2.40, p = .019, 95% CI [0.03, 0.38]. Memory-based fixation counts by presentation, response type, and similarity level are presented in Figure 2A.

**Figure 2.**
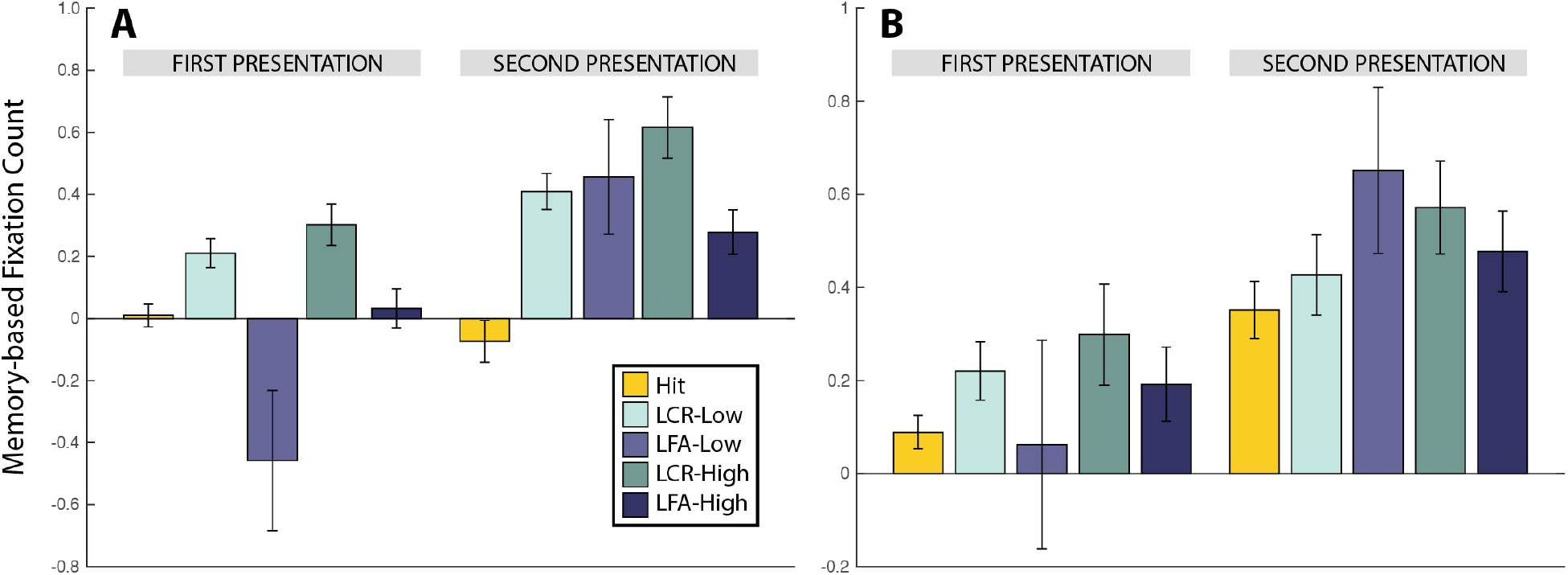
Memory-based fixation counts divided by presentation, outcome, and lure similarity. (A) In Experiment 1, fixation counts for subsequent LCRs were greater than for subsequent LFAs for both high- and low-similarity lures. LCRs also had a greater number of fixations than LFAs in Experiment 1. (B) In Experiment 2, there were more fixations during the second presentation than the first, but no other comparisons were significantly different. LCR = lure correct rejection, LFA = lure false alarm.

The more visually complex an image, the greater number of fixations on that image (Henderson, 2003). We attempted to eliminate the possibility of visual complexity as a confound by controlling for this in the present study. Effects that do not remain significant suggest that visual complexity was influencing fixation counts, not presentation or response type. To confirm this, we entered memory-based fixation counts into a linear mixed effects model with presentation number and outcome as fixed effects along with visual complexity scores. Presentation number and outcome were also included in the model as random slopes, with subject as a random intercept. We found that visual complexity significantly predicted fixation counts, β = 0.06, p < .001, 95% CI [0.04, 0.08] when other variables remained at baseline. Presentation number and response type, however, still significantly predicted fixation count when visual complexity was accounted for. Details from the model are included in Table 3.

**Table 3.**
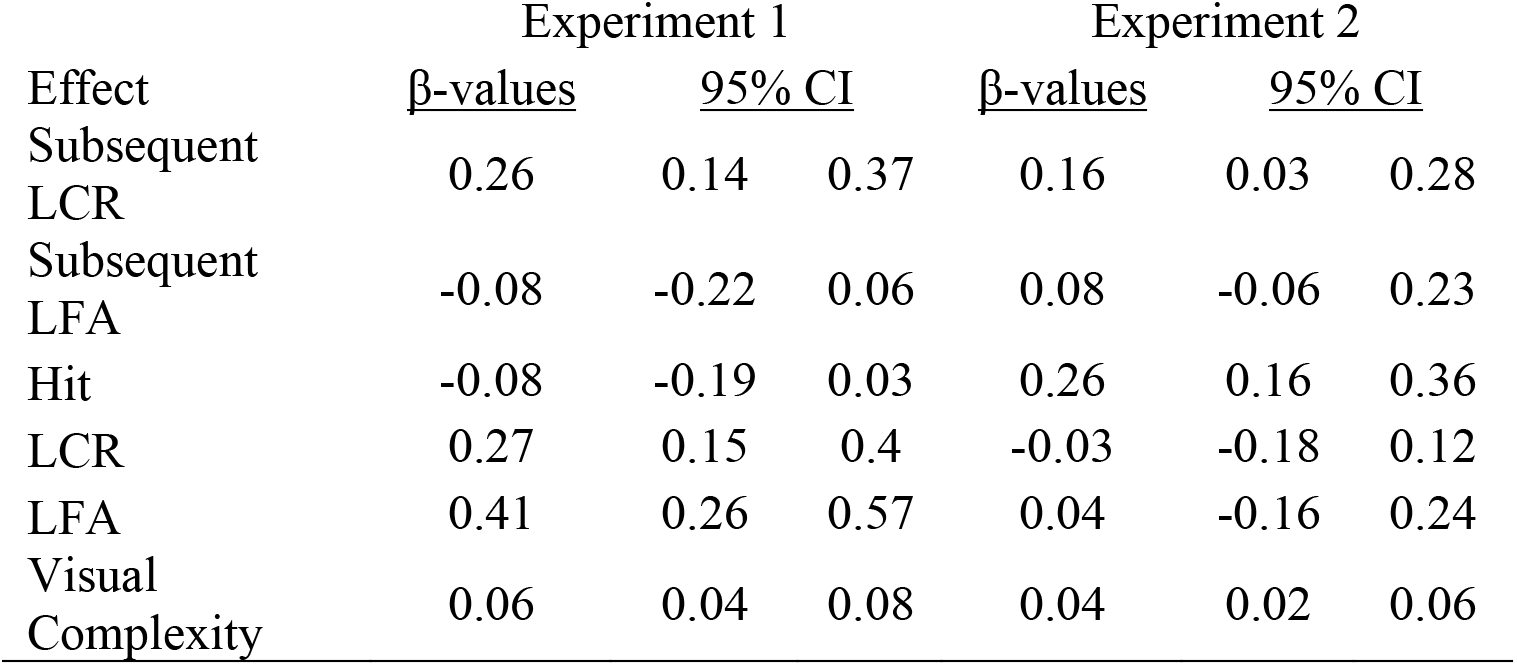
β-values and 95% confidence intervals for the mixed-effects regression models to investigate the effects of visual complexity on eye movements in Experiments 1 and 2. Note: Subsequent hits were used as the baseline in each model.

We next entered data for lure trials into a logistic mixed effects model with accuracy as the dependent variable (i.e., 1 for correct and 0 for incorrect). Raw fixation count at study, raw fixation count at test, and similarity level were fixed effects. We included all three as random slopes with the subject as a random intercept. Fixation counts were centered at the mean. After the model failed to converge, further examination revealed that some subjects had no or few false alarms to low-similarity lure trials. Accordingly, we removed the interactions with similarity level and also removed similarity level as a random slope. Successful convergence of the model showed that both fixations at study, β = 0.11, *p* < .001, 95% CI [0.06, 0.17], and fixations at test, β = 0.06, *p* = 0.02, 95% CI [0.01, 0.12], significantly predicted a correct response when similarity level was held to high-similarity trials, with each fixation increasing the odds of a correct response by 12.09% (95% CI [6.28%, 18.21%]) and 6.96% (95% CI [1.17%, 13.29%]), respectively. There was no significant interaction between fixations at study and fixations at test. We tested model fit by calculating a confusion matrix and found that 80.70% of the trials were correctly predicted.

### Experiment 2: Eye Movement Data

In Experiment 2, we found significant main effects of presentation, *F*(1, 34) = 26.47, *p* < .001, η_p_^2^ = .44, and response type, *F*(2, 68) = 3.65, *p* =.03, η_p_^2^ = .10 in the eye movement data but no significant interaction between presentation and response type, *F*(2, 68) = 0.04, *p* = .96, η_p_^2^ = .00. Post-hoc tests revealed significantly fewer fixations for hits than for LCRs, *t*(34) = −3.13,*p* < .01, 95% CI [−0.27, −0.06], and for LFAs, *t*(34) = −2.04,*p* < .05, 95% CI [−0.25, 0.00], though the upper limit of the 95% confidence interval very closely approached zero. We report no significant difference between fixation counts for LCRs and LFAs, *t*(34) =0.49, *p* = .63, 95 % CI [−0.11, 0.19]. Similar to Experiment 1 above, a linear mixed effects model revealed that visual complexity significantly predicted fixation counts, β = 0.04,*p* < .001, 95% CI [0.02, 0.06]. Parameters from the model are reported in Table 3.

We again examined the effects of similarity on memory-based fixations for lures using a repeated measures ANOVA with factors for presentation number (first, second), response type (LCR, LFA), and similarity rating (high, low). Similar to Experiment 1, we found a significant main effect for presentation, *F*(1,34) = 16.69, p < .001, η_p_^2^ = .33, though no other main effects or interactions were significant. Memory-based fixation counts by presentation, response type, and similarity are presented in Figure 2B.

We analyzed data for lure trials using the same logistic mixed effects model from Experiment 1. Fixations at study, β = 0.01, *p* = .70, 95% CI [−0.06, 0.08], and fixations at test, β = 0.004, *p* = .91, 95% CI [−0.07, 0.08], did not significantly predict correct responses, nor did the interaction between fixations at study and fixations at test, β = −0.02, *p* = .08, 95% CI [−0.05, 0.002]. We tested model fit with a confusion matrix and found that 78.08% of trials were correctly predicted.

### Experiment 2: MRI Data

We first identified regions involved in memory performance by contrasting the first and second presentations of correctly identified targets (i.e., the subsequent hit and hit conditions), yielding six clusters that showed repetition enhancement and eight clusters that showed repetition suppression, including clusters in bilateral hippocampal head (see Table 4, Figure 3). As our hypotheses concerned activation changes in the hippocampus, we focused all further analyses on the responses of the clusters identified in the hippocampal head bilaterally. In addition to the repetition suppression effect for hits used to define these clusters, we also found significant repetition suppression effects for high-similarity LCRs, low-similarity LCRs, and high-similarity LFAs in both left and right hippocampus.

**Table 4.** Characteristics of fMRI clusters of activation for repetition suppression (− direction) and repetition enhancement (+ direction) in the contrast of subsequent hits vs. hits. Coordinates are peak activation in MNI space. B=bilateral; L=left; R=right

| <u>Label</u> | <u>Direction</u> | <u>#Voxels</u> | <u>X</u> | <u>Y</u> | <u>Z</u> |
| --- | --- | --- | --- | --- | --- |
| B. Retrosplenial | - | 618 | -4 | -52 | 16 |
| L. Temporoparietal Junction | - | 551 | -58 | -68 | 23 |
| L. Hippocampal Head | - | 363 | -23 | -15 | -14 |
| R. Hippocampal Head | - | 347 | 21 | -14 | -14 |
| L. Middle Temporal Gyrus | - | 182 | -65 | -3 | -5 |
| L. Temporal Polar Cortex | - | 116 | -57 | 11 | -30 |
| L. Inferior Frontal Gyrus | - | 84 | -44 | 25 | -17 |
| L. Parahippocampal Cortex | - | 52 | -27 | -28 | -26 |
| B. Posterior Parietal | + | 5189 | 34 | -74 | 53 |
| B. Posterior Cingulate Cortex | + | 356 | 2 | -37 | 25 |
| R. Thalamus | + | 265 | 4 | -15 | 13 |
| R. Superior Temporal Gyrus | + | 246 | 69 | -45 | -3 |
| L. Thalamus | + | 203 | -12 | -17 | 9 |
| R. Cuneus | + | 114 | 12 | -72 | 11 |

**Figure 3.**
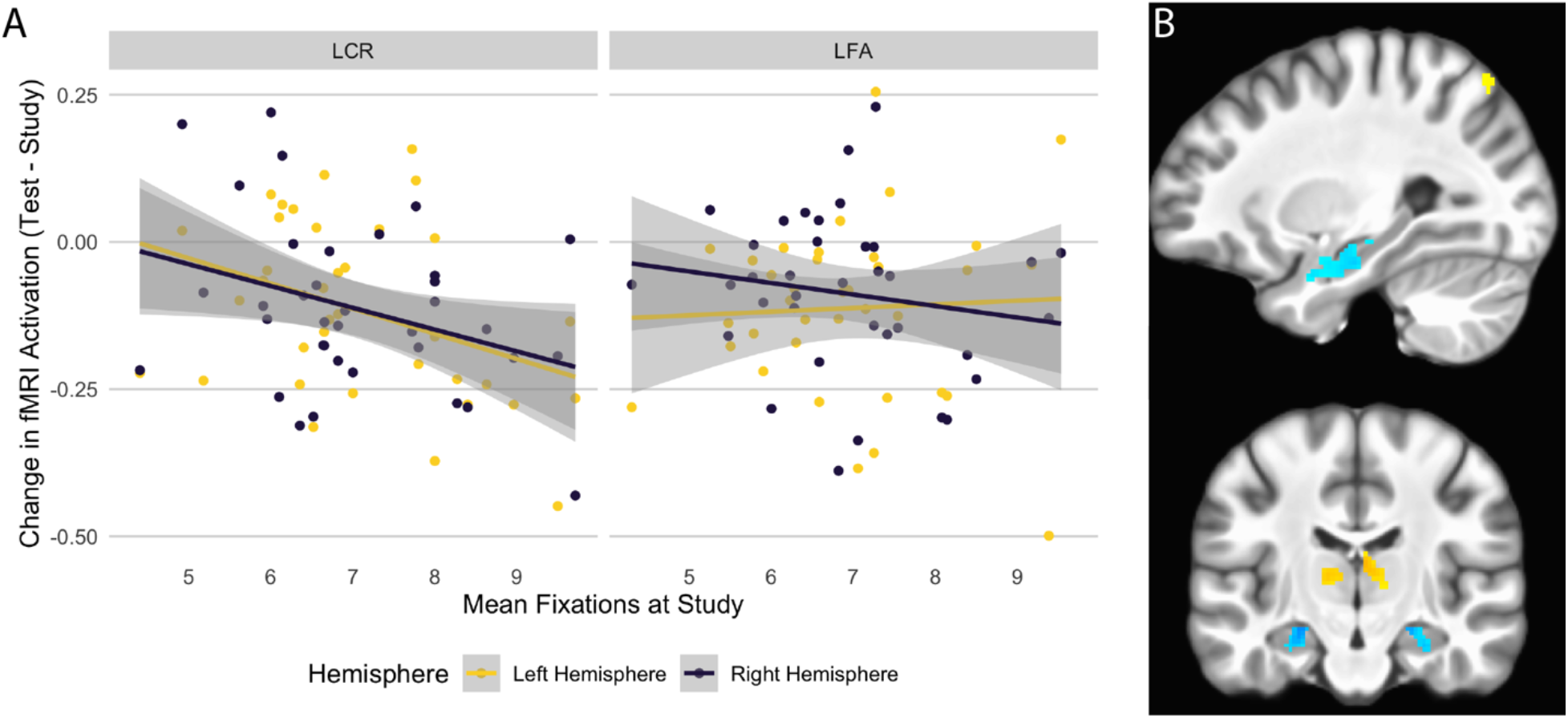
Fixations at study were significantly correlated with degree of repetition suppression for LCRs but not LFAs when controlling for target-lure similarity (A) in the left and right hippocampal head (B).

We next examined the relationship between fixations at encoding and hippocampal activation changes at test. As Experiment 1 demonstrated that target-lure similarity was a strong predictor of correct performance on lure trials and there were very few false alarms to low-similarity lures, we controlled for similarity by only examining activation for high-similarity lures. We examined the correlation between average number of fixations during encoding for high-similar lures and the average repetition suppression in the hippocampus (i.e., the difference in hippocampal activation between initial presentation of the target and subsequent presentation of the lure) for both LCRs and LFAs. We found a significant correlation between mean number of fixations at study and degree of repetition suppression for LCRs in left hippocampal head, *r* = −0.368, *t*(34) = −2.316, *p* = 0.027, and right hippocampal head, *r* = −0.336, *t*(34) = −2.069, *p* = 0.046. The correlation for LFAs was not significant in either left, *r* = 0.048, *t*(34) = 0.282, *p* = 0.780, or right hippocampal head *r* = −0.169, *t*(34) = −0.999, *p* = 0.325 (Figure 3A). The correlations were significantly different for the left hippocampus (z=-1.746, p=.04) but not the right hippocampus (z=-0.716, p=0.237). Taken together, these results indicate that eye movements at encoding were related to neural activation changes associated with successful, but not unsuccessful, subsequent mnemonic discrimination.

## Discussion

In two experiments, we examined the relative contributions of initial memory encoding strength and target-lure similarity on behavioral and neural indices of mnemonic discrimination. We replicated findings of previous research by demonstrating that eye movements at encoding predicted subsequent lure discrimination, emphasizing the importance of initial encoding strength for successful mnemonic discrimination (Molitor et al., 2014). Further, we found that eye movements at encoding also predicted subsequent repetition suppression in the hippocampus for correct lure discrimination. Replicating previous work (e.g., Kirwan and Stark, 2007; Motley and Kirwan, 2012), we found that target-lure similarity was a strong predictor of correct lure discrimination in the MST (internally replicated in Experiment 2). Further, the relationship between eye movements at encoding and subsequent repetition suppression in the hippocampus was strongest for high-similarity lure correct rejections and was much more modest for lure false alarms. Thus, while we demonstrate that initial encoding strength is important for subsequent mnemonic discrimination performance, behavioral performance is more strongly influenced by stimulus properties that are thought to drive pattern completion processes and encoding strength affects fMRI measures for correct mnemonic discrimination performance (i.e., pattern separation) rather than incorrect performance.

We found significantly more fixations at study for subsequent LCRs than for subsequent LFAs when we replicated the exact methods of Molitor et al. (2014) in Experiment 1. Logistic regressions for both Experiments 1 and 2 confirmed that fixations at study predicted the correct response of a trial at a greater rate than fixations at test, similar to interpretations from the previous study that highlight the importance of accurate memory traces for mnemonic discrimination (Molitor et al., 2014). However, in additional analyses we show that target-lure similarity level strongly associated with correct mnemonic discrimination performance, with more similar lures associated with more LFAs. These findings suggest that pattern completion is a major contributor to LFAs.

Experiment 2 did not reveal an interaction between presentation and response type that was expected based on the previous study and Experiment 1. This may have resulted from the change in task design, as the response period was fixed in Experiment 2 but self-paced in Experiment 1. Completion of the task in the scanner may have altered eye movement patterns as a result of performing in a changed environment. There were also fewer participants (about half as many) in Experiment 2 compared to Experiment 1. Power analysis from the first experiment required a sample size of 60 participants to achieve sufficient power, so it is possible that the analysis in Experiment 2 was underpowered given the common constraints of fMRI scanning on sample size (e.g., cost per scan and further restrictions on participant compatibility in the MRI environment). Future replications with the appropriate sample size are recommended to evaluate these potential accounts in greater detail.

Fixation counts can be influenced by many variables, and another possible explanation of a greater number of fixations for subsequent LCRs than for subsequent LFAs is differing levels of visual complexity. Fixation counts are greater when individuals are presented with stimuli that are more visually complex (Henderson, 2003). Both experiments showed a significant effect of visual complexity, with greater visual complexity resulting in a greater number of fixations to the image. However, the addition of visual complexity to the model did not change any of the significant results that the experiments found for effects of presentation, response type, or (when present) an interaction between the two, so it appears that significant differences between subsequent LFAs and subsequent LCRs cannot be explained by a difference in visual complexity between the target and the lure.

A growing body of literature has linked eye movements with hippocampal-dependent memory encoding. (Liu et al., 2017, 2018; Staudigl et al., 2017; Lucas et al., 2019; Kragel et al., 2020). Our fMRI findings are similar to those of Liu et al. (2017) who found that fixation count was related to the degree of hippocampal activation for novel, but not familiar faces and that the degree of repetition suppression for repeated stimuli was related to the number of fixations at encoding. Liu and colleagues used an incidental encoding paradigm where participants did not make overt memory judgments. Consequently, our results extend on the previous finding by demonstrating that the relationship between fixation count at encoding and the degree of subsequent repetition suppression in the hippocampus was specific to correct (LCR) but not incorrect (LFA) mnemonic discrimination judgments. Subsequent research has shown that the relationship between eye movements at encoding and hippocampal activation is attenuated in age (Liu et al., 2018). Given that mnemonic discrimination performance (Holden et al., 2012; Stark et al., 2013, 2015) and hippocampal responses (Reagh et al., 2018) are also attenuated with age, we suggest that the relationship between encoding-related eye movements and hippocampal activation in the MST would also be attenuated.

We found that eye movements at encoding were only correlated with repetition suppression in the hippocampus for LCRs but not LFAs. This finding is consistent with previous studies. For example, Liu et al. (2017) found a relationship between hippocampal activation and fixation count for novel but not familiar faces. While the previous study used an incidental encoding paradigm, it appears that the mnemonic discrimination task in the present study complicates the relationship between eye movements at encoding and repetition suppression in the hippocampus. If familiarity were the only factor influencing this relationship, then we would expect a correlation between fixations and repetition suppression for both LFAs and LCRs, as these trial outcomes result from participants indicating familiarity for either that exact stimulus or something like it, respectively. The observed pattern may be driven by either increased hippocampal pattern completion at retrieval for LFAs resulting in greater hippocampal activation (and thus no repetition suppression), or by differential encoding activity (increased for subsequent LCRs or decreased for subsequent LFAs). Further studies will be necessary to adjudicate these possibilities.

We did not observe differential responses across hippocampal subregions as might be predicted by computational models for pattern separation/pattern completion responses (e.g., Rolls and Kesner, 2006). Rather, we observed memory-related changes in functional activation in large clusters in the hippocampal head that did not differentiate between subregions. As we did not have strong predictions regarding the relationship between eye movements and activation differences in hippocampal subregions, we did not perform a priori region of interest analyses in hippocampal subregions. Future studies may wish to examine these effects in greater detail.

Target-lure similarity was the only predictor of accuracy that was significant across both experiments in the current study, supporting the idea that pattern completion drives false alarms to lure stimuli in the MST. Low-similarity lures were more likely to be correctly answered than high-similarity lures. When an explicit answer is required on a mnemonic discrimination task, a “recall to reject” strategy is commonly thought to be used (Hunsaker and Kesner, 2013). Such a strategy requires pattern completion because an individual must use the current image to recall the previously encountered image and compare them. If the overlap between the two images is too great, pattern completion may result in an “old” response to a lure (i.e., LFA).

One potential limitation to this study is the low number of false alarms to low-similarity lure trials through both experiments (mean 2.8 per subject, range 0-9). Accordingly, we controlled for similarity level by restricting analysis of the contributions of fixations at study and test to task performance to high-similarity lure trials. This restriction may have influenced the findings of our analyses. Such a limitation does point to the strong effect of lure pair similarity level, however, as participants did much better on the low-similarity lure trials than on high-similarity lure trials. Further research could alter the similarity level or stimulus presentation time to decrease performance slightly so that there is a better representation of false alarms in the low-similarity trials.

In summary, we demonstrate that in addition to target-lure similarity encoding-related eye movements predict subsequent mnemonic discrimination performance. Further, we demonstrate that when controlling for similarity, eye movements predict hippocampal memory responses for correct, but not incorrect mnemonic discrimination judgments, possibly due to enhanced encoding-related activation as indexed by eye movements. These findings contribute to a growing literature on the relationship between oculomotor and memory processes.

## Acknowledgments

We thank Torin Kelly, Elijah Möller, Emily Maxwell, Clara Rodriguez, Caitlyn Towne, Eric Brighton, Caroline Straw, Mia Felin, Ariana Hedges-Muncy, Jordan Clark, Athena Howell, Annelise Toolson, Cooper Hodges, Todd Winn, and Laurine Giraud-Carrier for help in data collection. Caitlyn Towne and Emily Maxwell were instrumental in programming the task for Experiment 2. Nathan Muncy offered consultation and advice on hippocampal segmentation and Ariana Hedges-Muncy answered many statistical inquiries. Dr. Steven Luke was also very helpful in providing feedback for analyzing the eye tracking data. This work was supported by a seed grant from the BYU MRI Research Facility.

